# Illuminating the Unconscious: Physical and Perceived Brightness in Cognitive Processing

**DOI:** 10.1101/2025.04.20.649724

**Authors:** Hirotaka Senda, Michael Makoto Martinsen, Hideki Tamura, Shigeki Nakauchi, Tetsuto Minami

**Affiliations:** Department of Computer Science and Engineering, Toyohashi University of Technology

**Keywords:** Continuous Flash Suppression, Illusion, Brightness Perception

## Abstract

This study investigated how physical luminance and perceived brightness affect breakthrough time (BT) under continuous flash suppression (CFS). Experiment 1 examined whether the glare illusion—which increases subjective brightness without altering actual luminance—would shorten BT compared to physically identical controls. The results revealed no difference, suggesting that subjective brightness alone does not expedite emergence into awareness. Experiment 2 assessed whether the partial suppression of the illusion’s inducers influenced detection speed and revealed that subjective brightness stimuli gained a BT advantage. Experiment 3 tested participants’ ability to discriminate real versus illusory brightness while stimuli remained suppressed; performance above chance for both conditions indicated that physical and perceived brightness cues were processed unconsciously. Together, these findings suggest that contextual brightness illusions are not simply lost below awareness—they can be discriminated in unconscious vision.

**Statement of Relevance:** Understanding how the visual system processes brightness illusions—even prior to awareness—is crucial for uncovering the brain’s deeper mechanisms of perception. This research clarifies the limits of unconscious visual processing by showing that illusions can be registered without consciousness but do not necessarily expedite emergence into awareness. These insights advance theoretical models of hierarchical vision and demonstrate that while early cortical areas prioritize actual luminance in determining whether a stimulus reaches awareness, higher areas can still encode illusory brightness beneath the threshold of consciousness. Thus, the study’s findings have broad implications for both basic neuroscience—refining how we think about the boundary between unconscious and conscious perception—and for applied fields seeking to harness or mitigate perceptual illusions.

**Research Transparency Statement:** *Preregistration:* All experiments were not preregistered.

*Conflicts of interest:* The authors declare that they have no competing interests.

*Data availability:* The code and data underlying the results presented in the study are available from the Open Science Framework repository (https://osf.io/u5der/).

*Ethics:* The Committee for Human Research of the Toyohashi University of Technology approved the experiments (2023-20). Written informed consent was obtained from all participants after the procedural details were explained to them.

*Funding:* This work was supported by JSPS KAKENHI (Grant Numbers JP22K17987 to H.T., JP24H01551 to T.M., JP23KK0183 to T.M., JP20H05956 to S.N.), JSPS Grant-in-Aid for JSPS Fellows (Grant Number JP24KJ1313), and Young Principal Investigator fund JPMJFS2121.

*Artificial intelligence:* During the preparation of this work, the authors used ChatGPT o1 and Grammarly to improve the language, and the manuscript was proofread by native English speakers through an English editing service. After using the tool and service, the authors reviewed and edited the content as needed, and they take full responsibility for the content of the publication.

## Introduction

While brightness is often linked to physical luminance, our perception of brightness is shaped by contextual and cognitive factors and frequently leads to striking mismatches between physical reality and subjective experience. A classic example is simultaneous brightness contrast (SBC), where identical gray patches appear brighter or darker depending on their background. Similarly, the glare effect dramatically alters perceived brightness: when a central area is surrounded by a radial luminance gradient, it appears significantly more luminous than it actually is (Agostini & Galmonte, 2002; Zavagno, 1999; Zavagno & Caputo, 2001). Studies have shown that this illusion can enhance perceived brightness by 20–35%, even with constant central luminance (Tamura, Nakauchi, & Koida, 2016; Yoshida, Mittner, Mantiuk, & Seidel, 2008), and offer powerful tools for studying how the visual system handles illusory brightness (Kinzuka, Sato, Minami, & Nakauchi, 2021; Suzuki, Minami, Laeng, & Nakauchi, 2019).

These illusions highlight that brightness perception is not driven solely by retinal input but is also shaped by contextual influences (Edward H. Adelson, 1993; Reid Jr & Shapley, 1988; Shapley & Reid, 1985; Zavagno, 1999). This suggests that both low-level mechanisms (Blakeslee & McCourt, 1999; Paradiso & Nakayama, 1991) and high-level interpretative processes (Edward H. Adelson, n.d.; Sinha et al., 2020; Williams, McCoy, & Purves, 1998) interact to construct brightness perception (Kingdom, 2003). A key question arises: if high-level inference contributes to illusions, such as the glare effect, does their perception require conscious awareness? While such illusions are well documented in conscious vision, it remains unclear whether similar effects occur under suppressed awareness.

Visual perception does not always equate to visual awareness, and many perceptual processes can occur outside conscious experience. Advances in unconscious experimental visual stimuli presentation—particularly continuous flash suppression (CFS)—have made it possible to render stimuli invisible while probing their neural processing. CFS presents dynamic noise to one eye and suppresses the target image in the other (Tsuchiya & Koch, 2005). Stimuli can remain suppressed for seconds before awareness is reached. The breakthrough time (BT) quantitatively measures unconscious processing efficiency: shorter BTs indicate more efficient processing. This breaking-CFS paradigm has been widely used to explore how different stimuli are handled without awareness (Jiang, Costello, & He, 2007; Song & Yao, 2016; Stein, Hebart, & Sterzer, 2011). Prior studies have shown that physical features, such as luminance and contrast, influence suppression duration. For example, Song and Yao (2016) reported that increasing a stimulus’s luminance improved discrimination accuracy—even when subjects reported not seeing it—which indicates that stronger signals are prioritized by the visual system. However, it remains unclear whether perceived brightness from illusions, such as the glare effect, offers the same advantage. If illusory brightness shortens BT, it would suggest that subjective perceptual qualities, not just objective physical properties, can facilitate unconscious processing. This would extend the evidence that unconscious vision integrates complex and context-dependent information (Mudrik, Breska, Lamy, & Deouell, 2011; Stein et al., 2011). For example, suppressed stimuli can still show object–context integration (Mudrik et al., 2011), some illusions persist in the absence of awareness of the inducing context. Harris, Schwarzkopf, Song, Bahrami, and Rees (2011) reported that SBC illusions occur even when their surroundings are rendered invisible, which suggests that brightness-related contextual computations can operate unconsciously. If the glare illusion also influences BT under suppression, it would further support the idea that unconscious visual processing encompasses not only low-level features, such as luminance, but also higher-order perceptual constructs, such as induced brightness.

Given these open questions, we present multiple experiments using the glare illusion to examine how physical and perceived brightness interact under suppression. The glare illusion was selected due to its robust and well-documented effects on perceived brightness (Tamura et al., 2016; Zavagno, 1999). Collectively, these experiments bridge the research on brightness perception and visual awareness and offer insight into the extent to which brightness enhancements depend on conscious vision.

## Experiment 1

Experiment 1 tests whether glare-induced brightness enhancements reduce BT compared to physically identical, non-illusory targets to directly examine whether subjective brightness facilitates unconscious access.

### Participants

A total of 24 healthy volunteers (4 females and 20 males; average age: 21.38 ± 1.32) with normal or corrected-to-normal vision from Toyohashi University of Technology participated in the experiment. The sample size was determined with G*power 3.1 (Faul, Erdfelder, Lang, & Buchner, 2007) to achieve a medium effect size in a two-tailed t-test with □ = *0.05* and □ = *0.80*. The experiment was conducted with the approval of the Ethics Committee for Human Research at Toyohashi University of Technology and adhered strictly to the approved guidelines of the committee and the Declaration of Helsinki. Written informed consent was obtained from the participants after the experimental procedures were explained. This study was not preregistered.

### Apparatus

The experiment was conducted in a dark room, with a chin rest installed to maintain a distance of 80 cm from an LCD monitor (VIEWPixx/3D, VPixx Technologies Inc) with a resolution of *1920* × *1080* pixels and a refresh rate of 120 Hz. Prior to the experiment, the monitor was calibrated using a spectroradiometer (SR-3AR; Topcon, Tokyo, Japan) to a white point of D65 and a gamma of 1 to ensure a linear relationship between the RGB values and the brightness output which is crucial for the linear gradient of the glare illusion. The stimuli were presented using MATLAB R2023a and Psychtoolbox-3 (Brainard & Vision, 1997; Mario et al., 2007; Pelli & Vision, 1997). Participants wore active shutter 3D glasses (3D Vision Wireless, NVIDIA) to present the target stimulus and mask stimuli to different eyes during the experiment.

### Target and masking stimuli

Four target stimuli were specifically designed and presented in this experiment, as shown in Figure 1 (b). The stimulus on the far left is an example of a glare illusion, combining small circles arranged in a circular pattern and filled with a linear gradient changing from black to white (Suzuki et al., 2019). To test the robustness of the glare illusion across luminance polarities, we created both white- and black-centered versions of the circular glare patterns described by Tamura et al. (2016). Their findings revealed brightness enhancement over a wide luminance range, including conditions where the central region was darker than the background, and suggested that the effect is not limited to self-luminous appearances. This aligns with the work of Agostini and Galmonte (2002), who demonstrated that luminance gradients—whether from dark to light or vice versa—can strongly modulate achromatic simultaneous contrast. We included black glare stimuli based on these results to examine whether the illusion persists and maintains its perceptual strength under reversed contrast polarity. As control stimuli, we created stimuli that do not produce a change in the sensation of brightness Specifically, the gradient was rotated by 45 degrees (bottom left, bottom right) while the same number of black and white pixels was maintained as the glare illusion stimuli. For all four target stimuli, the brightest RGB value was set to 255 (*90* cd/m^2^), and the darkest RGB value was set to 0 (*0.078* cd/m^2^). The gradient had a linear profile, where the RGB value was modulated from zero to 255, and the size of each target stimulus was set to *3.5*^°^ × *3.5*^°^. For the CFS experiment, masking stimuli were designed to suppress each target stimulus. The mask stimuli were created using MATLAB R2023a by randomly overlaying multiple circles of size *1.0*^°^ ± *0.25*^°^ onto a canvas of *9.5*^°^ × *9.5*^°^ (Nuutinen, Mustonen, & Häkkinen, 2018). Owing to the random luminance values of each generated circular mask stimulus image, luminance equalization among the circular mask images was performed using the SHINE toolbox (Willenbockel et al., 2010). During the experiment, a black frame (*10.9*^°^ × *10.9*^°^) was presented to facilitate binocular fusion within which the target and mask stimuli were displayed.

**Figure 1.**
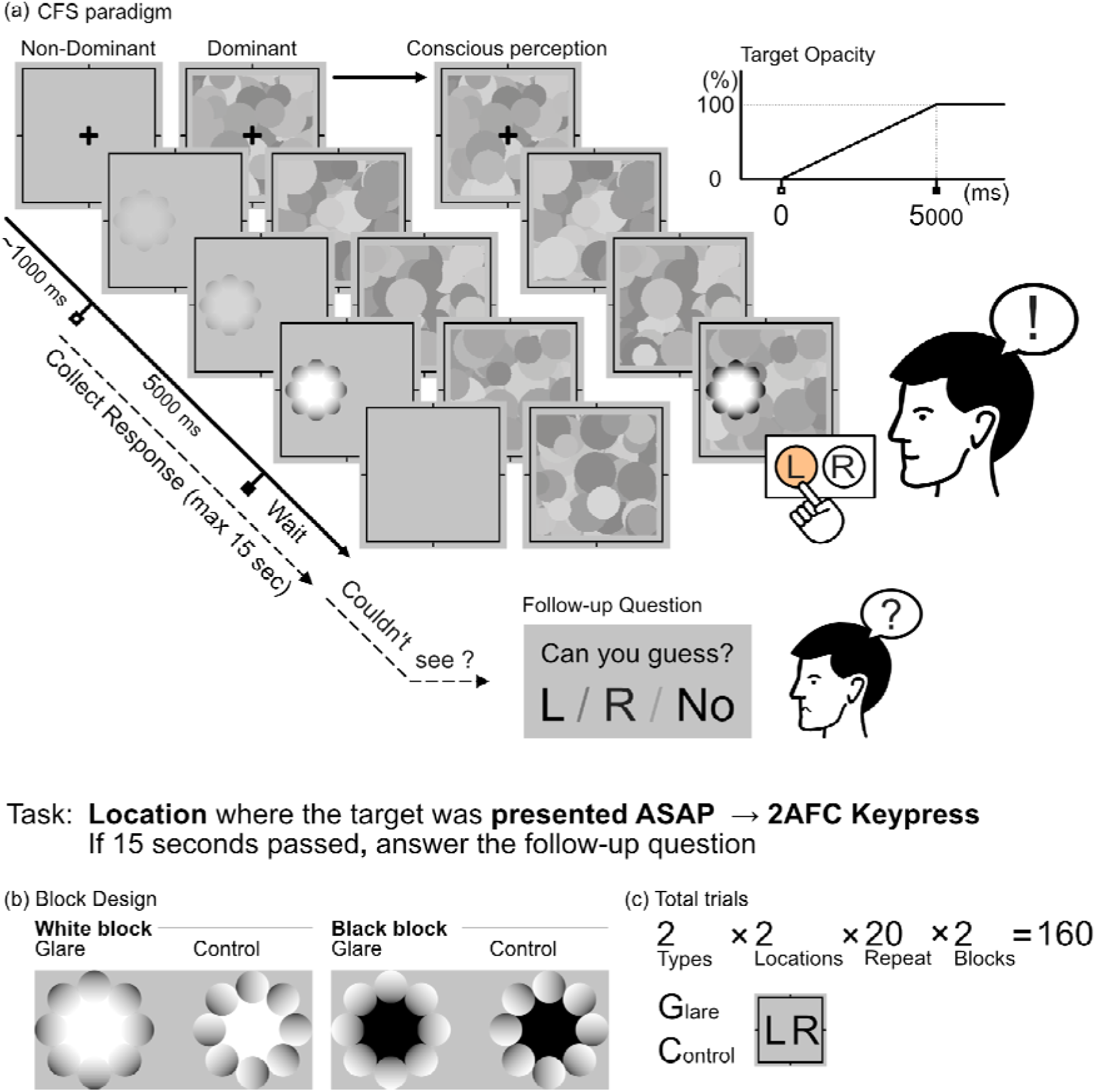
Experiment 1 paradigm. (a) The flow of the experiment. (b) Types of target stimuli used in each block (c) The total number of trials for the entire experiment.

### Procedure

The participants sat in front of the monitor with a chin rest and viewed the stimuli directly from the front while wearing 3D shutter glasses. Prior to the main experiment, the participants performed a CFS dominant-eye test to determine which eye was more susceptible to suppression, as one eye typically exhibits stronger suppression than the other. The method involves randomly presenting an arrow to either the left or right eye with decreasing transparency and increasing contrast, while presenting a mask stimulus to the non-arrow viewing eye. Participants responded with the corresponding key indicating the direction the arrow was pointing when it became visible, and the eye with the faster average BT was determined to be the dominant eye (Yang, Blake, & McDonald, 2010).

The main experiment was conducted as follows (Figure 1(a)). A target stimulus with 100% transparency was presented to the dominant eye on either the left or right side of the screen. The transparency of the target stimulus was then decreased by 5 seconds until it became opaque. Participants were instructed to press the left or right key as soon as the stimulus became visible, corresponding to the position of the target stimulus. Throughout the entire trial, masking stimuli were presented to the dominant eye at a frequency of 10 Hz and remained onscreen until a key was pressed. If the target stimulus was not visible even after 15 seconds, stimulus presentation for both eyes was halted, and the participants were instructed to guess where they thought the target stimulus had been presented.

Each participant completed 160 trials. The trials were organized into two blocks to compare the Glare and Control conditions; specifically, one block used only the White stimuli, and the other used only the Black stimuli. Each block consisted of 80 trials (2 conditions of Glare/Control × 2 stimulus presentation locations × 20 repetitions), and the presentation order was counterbalanced within the blocks (Figure 1(b,c)). Participants were not provided with any prior knowledge regarding the target stimuli.

### Analysis

Twenty-two participants were included in the analysis. One participant who did not perceive suppression adequately under CFS and another participant with a task accuracy rate of less than 70% were excluded from the analysis (Martinsen, KINZUKA, SATO, MINAMI, & NAKAUCHI, 2023; Martinsen et al., 2024).

In Experiment 1, trials where the BT was less than 100 ms or more than 15 seconds and trials where the participant responded incorrectly to the quadrant in which the stimulus appeared were excluded from the analysis. Subsequently, outliers in BT were removed for each participant using the 3□ (□ : standard deviation) rule and using R statistical software (version 4.3.2). On average, 95.3% of the trials per participant were retained for analysis. In 1.99% of the trials, the participants reached the follow-up question and answered incorrectly, indicating that breakthrough suppression was not achieved.

A Shapiro Wilk test was conducted to assess the normality of the data. The results for the White condition approximated a normal distribution (white-glare: □= *0.99*, □= *.977*; white-control: □= *0.94*, □= *.207*), whereas the results for the Black condition did not (black-glare: □= *0.87*, □= *.007*; black-control: □= *0.98*, □= *.857*). Therefore, a t-test was used to compare the BTs between the Glare and Control conditions in the White block. In contrast, a Wilcoxon signed-rank exact test was used for the same comparison in the Black block due to non-normality.

### Results

The experimental results for each stimulus are shown in Figure 2, where the vertical axis represents the average BT for each condition, and the horizontal axis indicates the Glare/Control conditions. A t-test was conducted to compare the BTs between the Glare and Control conditions in the White block (white-glare: □= *3.96*, □□= *0.35*, white-control: = *4.22*, = *0.34*), which showed no significant difference (□_□_ = *-0.23*, 95% CI [*-0.63,0.17*], □(*21*)= *-1.20*, , Cohen’s d = -0.149). In addition, a Wilcoxon signed-rank test was conducted to compare the BT between the Glare and Control conditions in the Black block (black-glare: , , black-control: , ), which also showed no significant difference ( , , r = 0.023). These findings indicate that category differences did not affect suppression in CFS.

**Figure 2.**
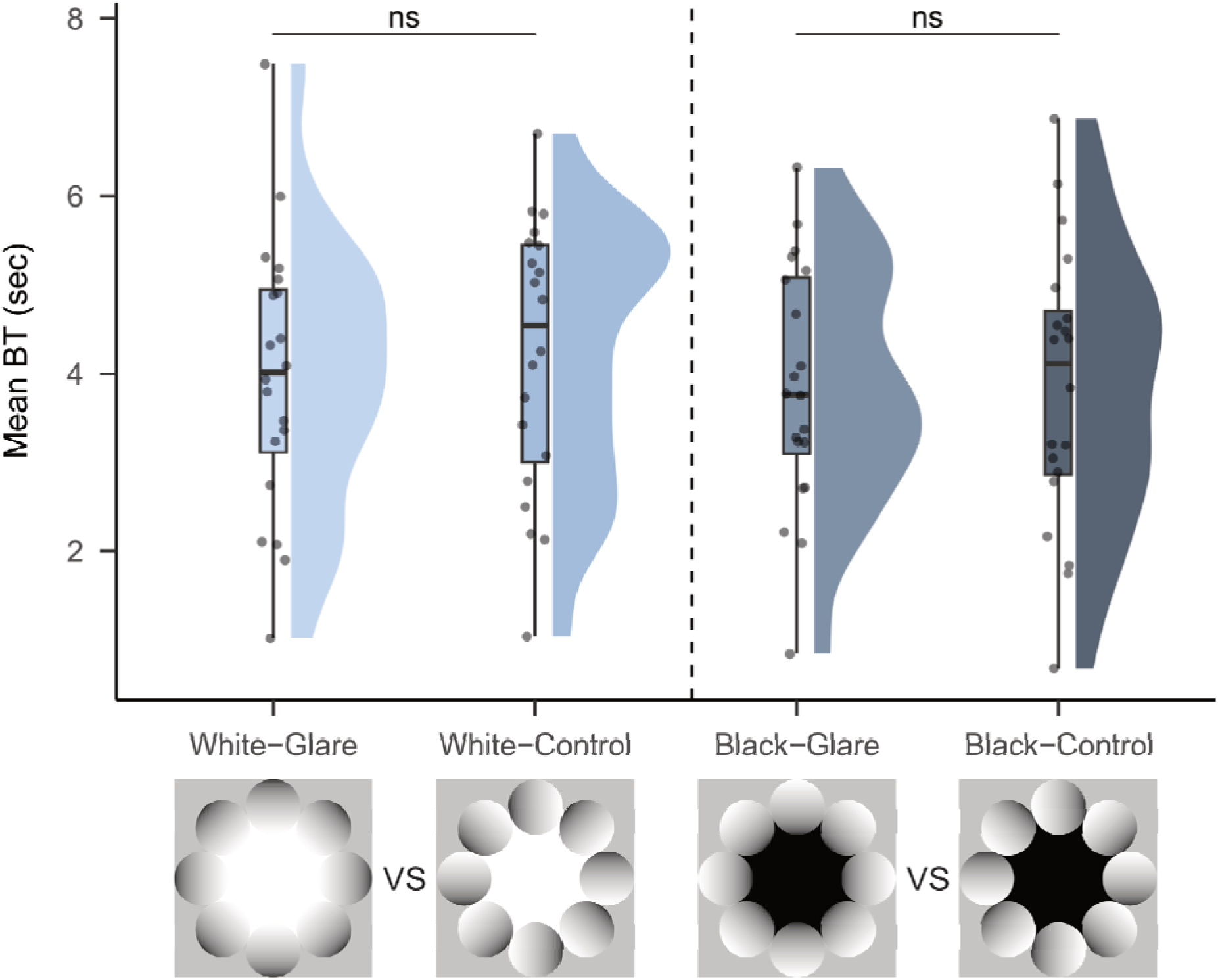
Raincloud plots showing the mean durations of CFS suppression across conditions in Experiment 1. Each plot displays individual data points (dots), the probability density of the distribution (cloud), and a boxplot showing the median and interquartile range. The white glare and white control conditions are shown side by side.

In Experiment 1, no significant differences were observed between the glare and control stimuli under either the white or black conditions. This result suggests that when the entire stimulus is suppressed from conscious awareness, the brightness illusion may not be processed. However, we considered that some components of the stimulus—particularly the inducers— might still undergo unconscious processing even when the global percept does not emerge. If the brightness illusion arises from the local processing of the inducers (i.e., the radial or linear gradients), then the suppression of only the central region while the inducers are allowed to remain visible could still lead to unconscious modulation. On the other hand, if the illusion requires the global integration of the entire stimulus, including the central region, then the removal of the inducer should eliminate any unconscious effect. Based on this reasoning, Experiment 2 was designed to test whether the suppression of only the inducers of the glare illusion would influence perceptual or behavioral outcomes, which allowed us to isolate the contribution of local versus global processing in unconscious brightness perception.

## Experiment 2

In Experiment 2, we isolated the glare illusion’s inducer gradient to assess whether partial suppression of the gradient altered how quickly the illusion became visible—probing the respective contributions of physical and contextual cues.

### Participants

A total of 24 healthy volunteers (6 females and 18 males; average age: 22.50 ± 0.87 years) with normal or corrected-to-normal vision participated in the experiment as in experiment 1. The experiment was conducted with the approval of the Ethics Committee for Human Research at Toyohashi University of Technology and adhered strictly to the approved guidelines of the committee and the Declaration of Helsinki. Written informed consent was obtained from the participants after the experimental procedures were explained. This study was not preregistered.

### Target and masking stimuli

In Experiment 2, we used the same white stimuli as those used in Experiment 1 (white-glare, white-glare in 3(a)) while modifying only the masking stimuli. Specifically, two white circles (RGB 255, *90* cd/m^2^) with a size *1.5*^°^ × *1.5*^°^ were overlaid *2.0*^°^ away from the center of the masking stimuli to mimic a “hole” in the mask, and participants were able to perceptually observe the center of the target during the entire paradigm while only the inducer was suppressed from consciousness (Figure 3(b)).

**Figure 3.**
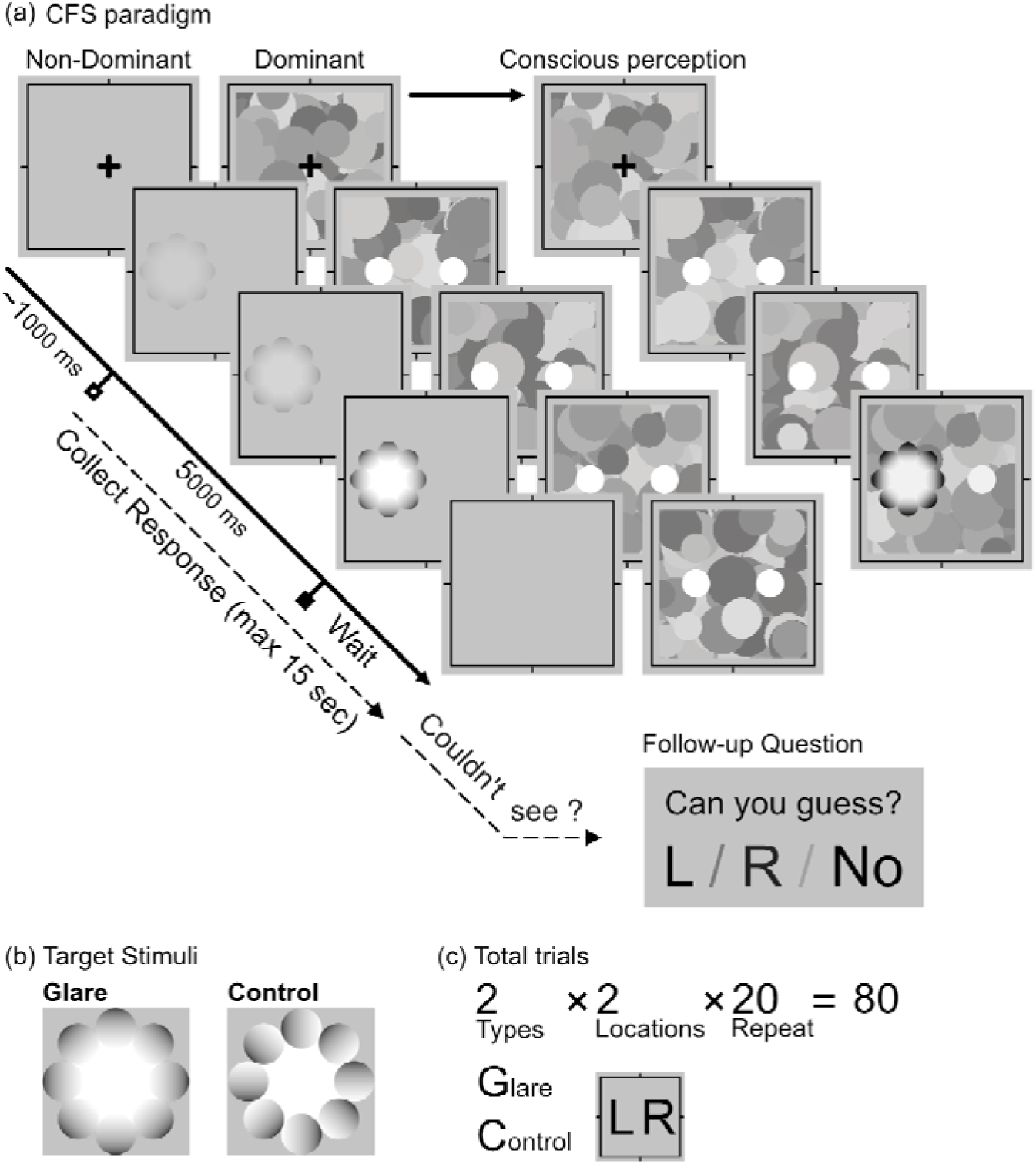
Procedure for Experiment 2. A specialized mask targeted only the glare inducers, leaving the central region visible under continuous flash suppression. This design tested whether the selective suppression of the gradients responsible for the glare illusion would alter detection speed or the perceptual impact of the illusion.

### Procedure

The flow of Experiment 2 was identical to that of Experiment 1; a fixation was presented for 1 second, after which the transparency of the target stimulus started decreasing over 5 seconds. Participants performed a detection task by pressing the corresponding key (L or R) on the keypad to indicate where the target stimuli appeared on the screen. If a participant did not press the keypad within 15 seconds, a follow-up question was presented to determine whether the participant had seen the target stimuli at the last moment. The flow of the experiment is shown in Figure 3(a).

Each participant completed 80 trials (Glare/Control × 2 presentation locations × 20 repetitions), and the presentation order was counterbalanced within the blocks (Figure 3(b,c)). As in Experiment 1, participants were not given any prior information about the target stimuli.

### Analysis

Twenty-one participants were included in the analysis. One participant with poor performance (fewer than 70% correct trials) and two participants whose mean BTs were identified as significant outliers were excluded. On average, 96.3% of the trials per participant were included in the analysis. In 2.8% of the trials, participants reached the follow-up question and failed to report the stimulus location correctly, indicating a failure to achieve breakthrough suppression. A Shapiro Wilk test was conducted to assess the normality of the data. The results for both the Control and Glare conditions approximated a normal distribution (Control: □= *0.95*, □= *.376*; Glare: □= *0.93*, □= *.112*). Therefore, a t-test was performed to compare BTs between the Glare and Control groups.

### Results

The results are shown in Figure 4, where the vertical axis represents the average BT for each condition, and the horizontal axis indicates the Glare/Control conditions. The t-test conducted between the two conditions (Glare: □= *5.17*, □□= *0.35*, Control: □= *4.53*, □□= *0.25*) revealed a significant difference (□_□_ = *0.64*, 95% CI [*0.20,1.07*], □(*20*)= *3.04*, □= *.006*, Cohen’s d = 0.457), which indicated that the BT for the glare illusion was significantly faster than that for the control stimuli.

**Figure 4.**
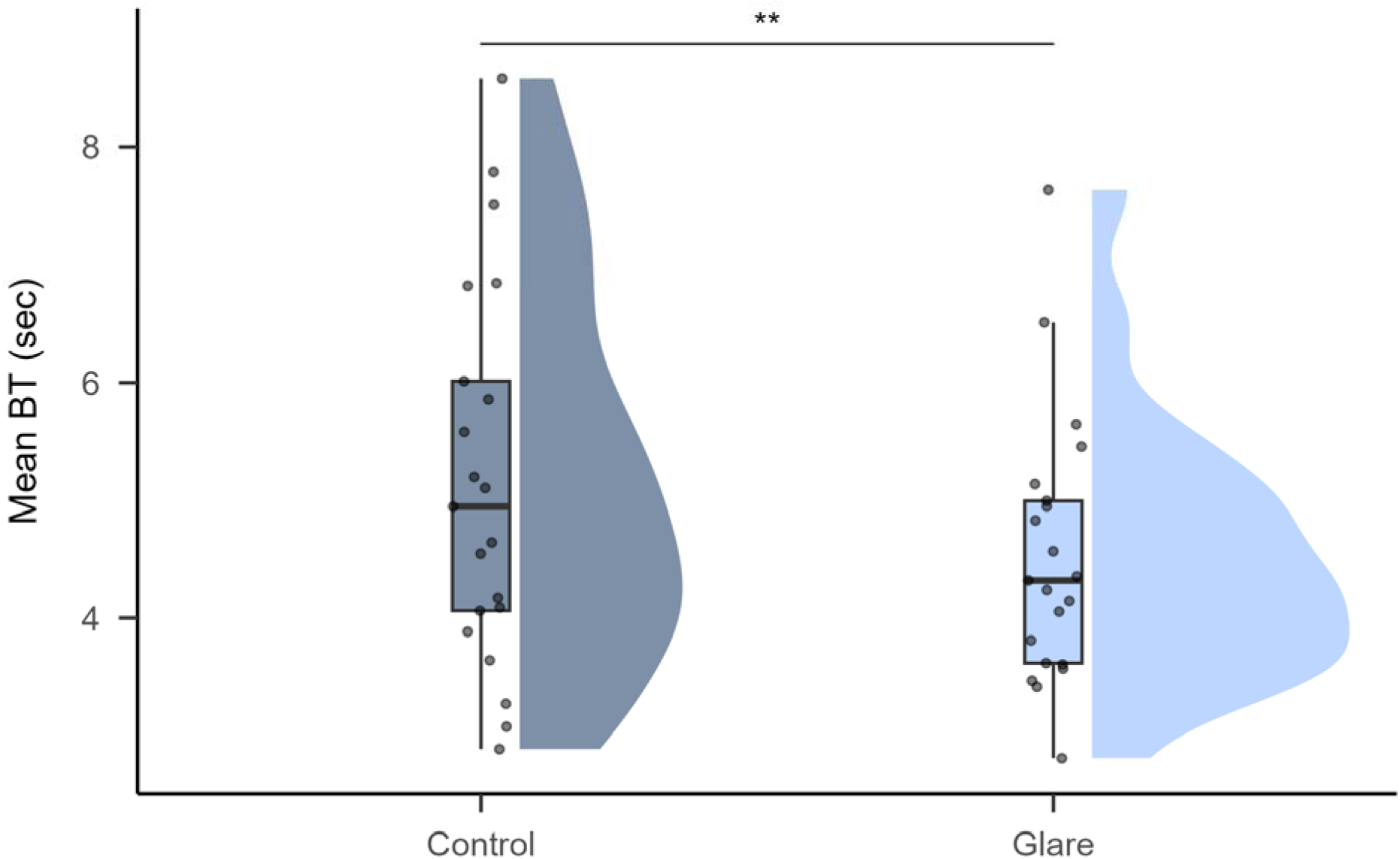
Results of Experiment 2. The vertical axis represents the average BT in which only the inducers were suppressed. Each gray dot indicates the participant’s mean BT.

Our results suggest that the glare illusion influences unconscious processing and that physical luminance and perceived brightness can affect the speed of visual breakthroughs to consciousness. However, these results could be explained by differences in stimulus complexity between the two conditions rather than the brightness illusion itself (Song & Yao, 2016). To address this, we conducted an additional experiment in which the target and control stimuli were presented side by side. Participants were instructed to select the brighter stimulus, which enabled us to assess whether the brightness illusion, which enhances perceived brightness without increasing luminance, would be preferentially chosen even when participants were unaware of the target stimulus.

## Experiment 3

In Experiment 3, illusory brightness enhancements were compared with actual luminance increases to determine whether observers detected them similarly under suppression, which shed light on how deeply illusory signals are processed without awareness (Harris et al., 2011).

### Participants

A total of 19 healthy volunteers (18 males and 1 female; average age: 22.42 ± 1.18) with normal or corrected-to-normal vision participated. The sample size was determined with G*power 3.1 (Faul et al., 2007) to achieve a medium effect size in a one-way t-test with □= *0.05* and □= *0.80*. The experiment was conducted with the approval of the Ethics Committee for Human Research at Toyohashi University of Technology and adhered strictly to the approved guidelines of the committee and the Declaration of Helsinki. Written informed consent was obtained from the participants after the experimental procedures were explained. This study was not preregistered.

### Target and masking stimuli

In Experiment 3, to investigate whether the effect of increased sensation of brightness due to the glare illusion is detectable even unconsciously, stimuli with the glare illusion and halo illusion placed on the left and right were used, as were stimuli with circles of different luminance levels placed on the left and right. Figure 6(b) describes the two sets of target stimuli: Real and Illusion conditions. The Real condition presented circles with Y values of *90* cd/m^2^ and *64* cd/m^2^. The Illusion condition presented an illusion image with a central area Y value of *64* cd/m^2^. The stimuli for the Real condition and the Illusion condition are shown in Figure 5. The Y values of these target stimuli were set to reflect a 40% difference in physical luminance, as Tamura et al. (2016) reported that the glare illusion can increase the sensation of brightness by (Tamura et al., 2016). For masking stimuli, circular mask stimuli identical to those used in Experiment 2 were employed. The presentation locations of the target stimuli were filled with matching colors to prevent partial suppression. As in Experiment 2, a black frame was presented for binocular fusion, inside which the mask and target stimuli were displayed.

**Figure 5.**
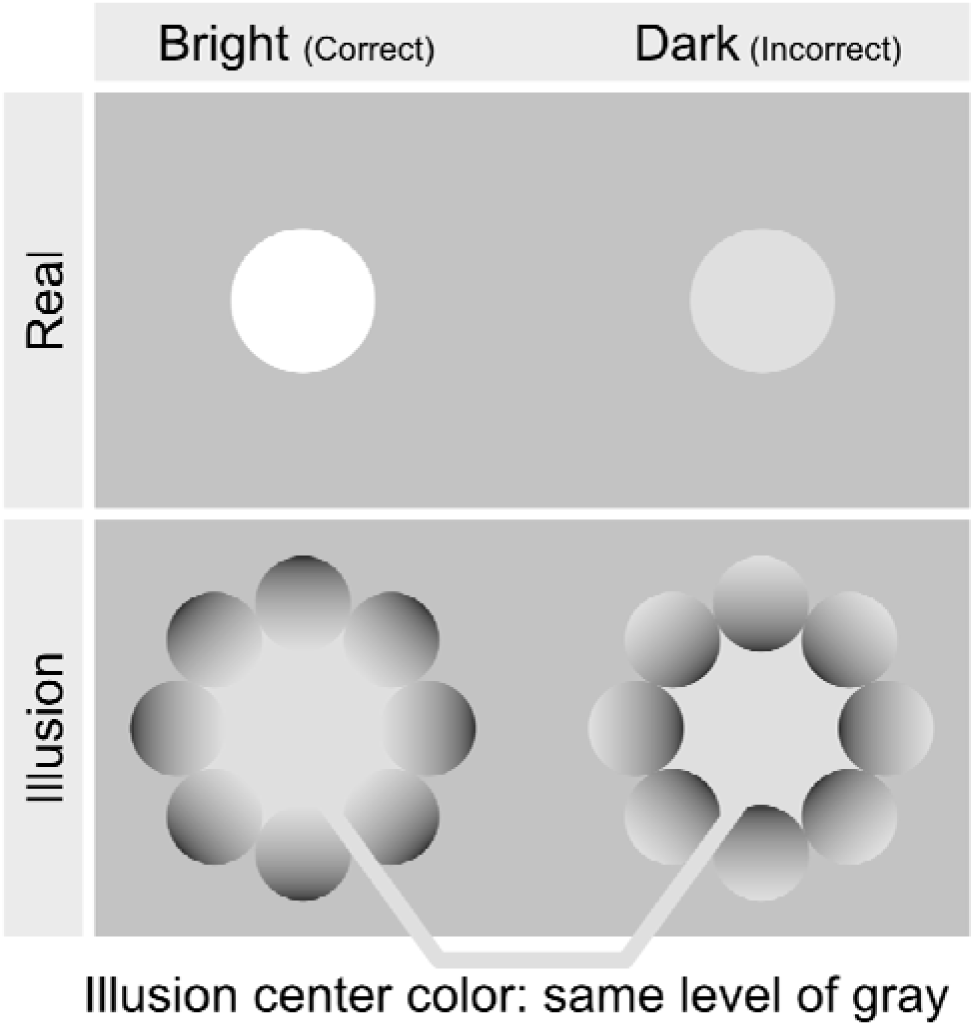
Target stimuli used in Experiment 3. In the Real condition, a pair of circles with different physical luminance (bright vs. dark) was presented. In the Illusion condition, glare and halo stimuli with identical central luminance were presented.

**Figure 6.**
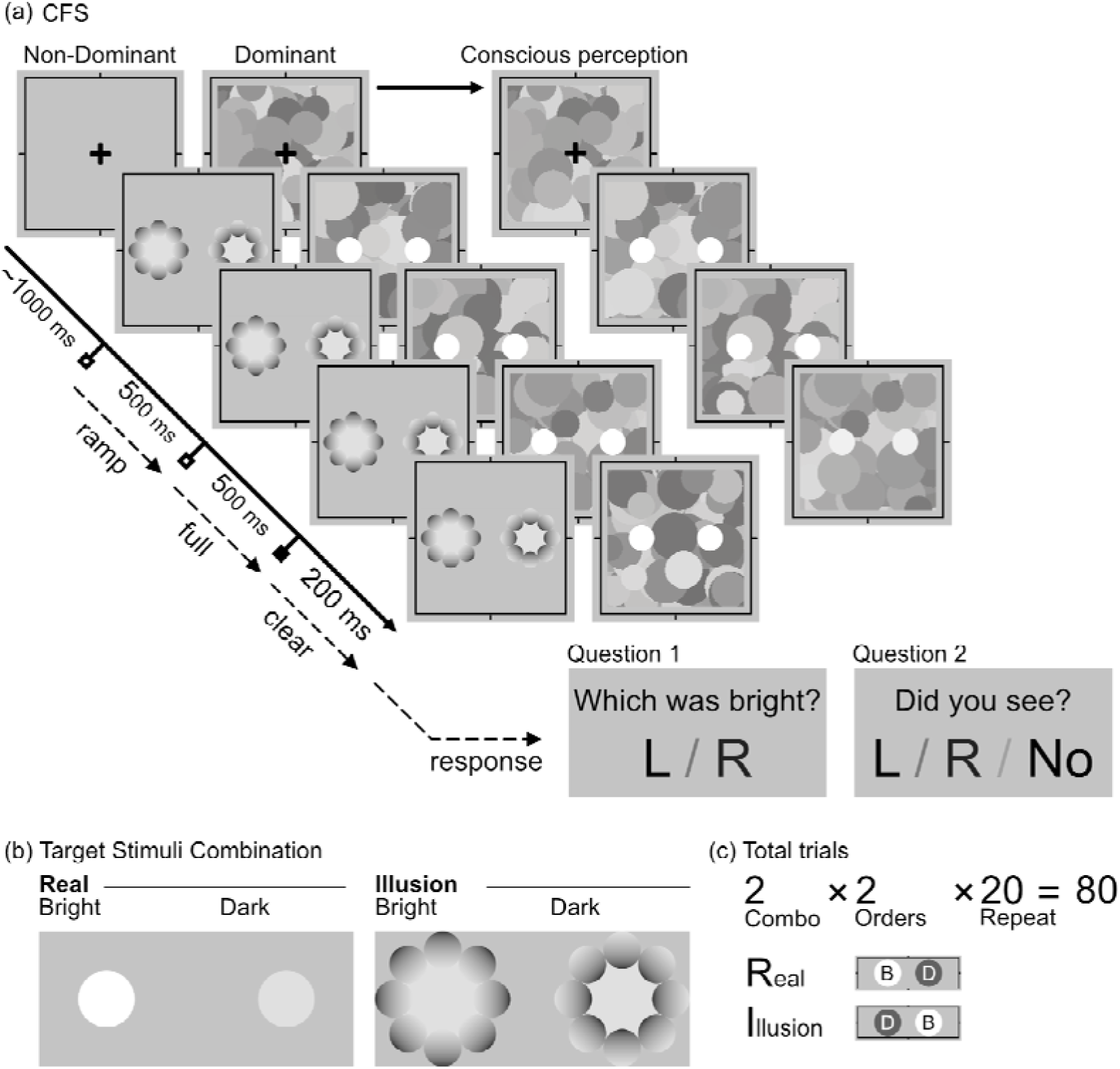
Procedure of Experiment 3. In each trial, one stimulus was physically brighter, while the other was made brighter via a glare illusion or physically, yet both remained suppressed from a conscious view. Participants indicated which side appeared brighter, which allowed the assessment of whether illusory brightness could be detected unconsciously.

### Procedure

The flow of Experiment 3 was as follows: First, a fixation point was presented for 1 second, followed by the presentation of the mask stimulus to the dominant eye and a target stimulus with 100% transparency to the non-dominant eye. The transparency of the target stimulus gradually decreased over 0.5 seconds. In accordance with the research of Harris et al. (2011), to prevent aftereffects, a mask stimulus was presented to both the dominant and non-dominant eyes for 0.2 seconds after the target stimulus presentation (Harris et al., 2011). After the stimuli were presented, the participants performed a brightness response task in which they compared the stimuli presented on the left and right sides and responded by pressing the corresponding key to indicate which one appeared brighter, regardless of visibility. In this task, selecting the brighter circle was considered correct in the Real condition, and selecting the glare illusion was considered correct in the Illusion condition. Finally, the participants were asked whether they could see the target stimulus. The flow of the experiment is shown in Figure 6.

### Analysis

To investigate the effect of illusions under unconscious conditions, trials in which participants responded that they had seen the target stimulus were excluded from the analysis. Additionally, one participant with poor suppression performance (visibility over 70%) was excluded, which resulted in a final sample size of 18 participants.

For trials in which participants reported not seeing the target stimulus, we calculated the accuracy rate of the brightness response task separately for the Real and Illusion conditions. We then tested whether this accuracy rate was significantly higher than the chance level. On average, 84.6% of the trials per participant were successfully suppressed from awareness. In addition, 70.6% of the trials per participant were utilized in the analysis.

A Shapiro□Wilk test was conducted to assess the normality of the data. The Real condition did not approximate a normal distribution, whereas the Illusion condition did (real: □= *0.38*, □< *.001*; illusory: □= *0.95*, □= *.392*). Therefore, a one-sample t-test was performed for the Illusion condition, while a Wilcoxon signed–rank exact test was used for the Real condition due to non-normality.

### Results

The results are shown in Figure 7, where the vertical axis represents the average discrimination rate for each condition, and the horizontal axis indicates the Real/Control conditions. The analysis focused on participants’ ability to discriminate differences in perceived brightness, with visual imagery employed to support the findings. The Wilcoxon signed-rank test suggested that the discrimination rate was significant compared to 50% (□ = *171.00*, □< *.001*, r = *0.909*), which indicated that participants were able to discriminate physically brighter stimuli from the gray stimuli as anticipated. Furthermore, the one-sample t-test revealed that glare stimuli were correctly selected above the chance level, which was set at 50% (□= *0.70*, 95% CI [*0.61,0.79*], □(*17*)= *4.80*, □< *.001*, Cohen’s d = 1.132).

**Figure 7.**
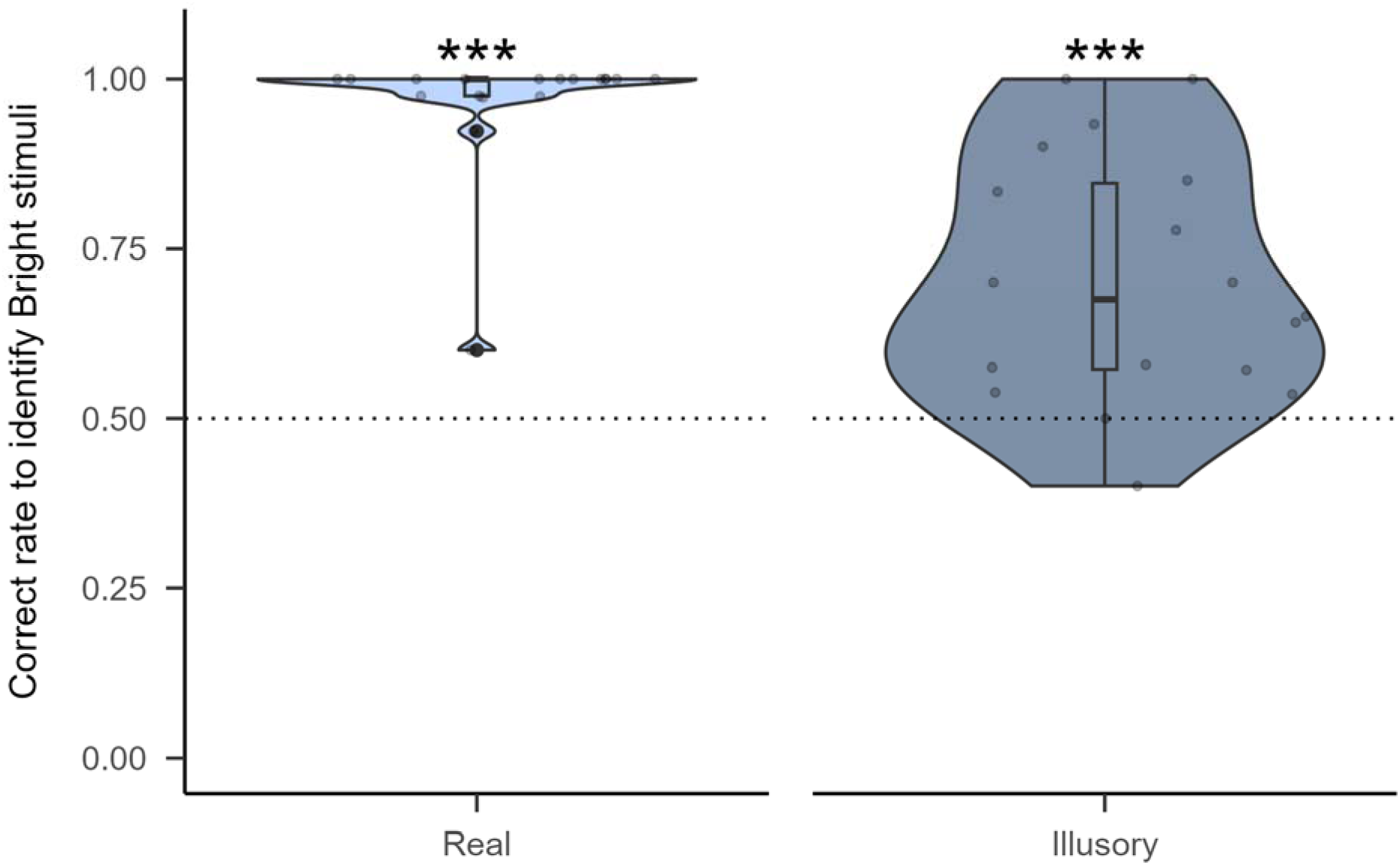
Results of Experiment 3. The vertical axis represents the accuracy of discrimination between physically brighter stimuli and those made brighter by a glare illusion while both remained suppressed. Each gray dot indicates the participant’s mean accuracy.

## General Discussion

This research examined how physical luminance and illusory brightness influence unconscious visual processing under CFS. Experiment 1 tested whether the glare illusion—which subjectively enhances brightness—would shorten the BT of suppressed stimuli. Contrary to expectations, illusory brightness did not expedite access to awareness, which suggested that early unconscious mechanisms prioritize actual luminance over perceptually amplified brightness (Song & Yao, 2016). Experiment 2 shifted the focus to the inducer gradients underlying the glare illusion. While these gradients did not reduce BT either, participants could still detect the presence of the illusion without conscious awareness, which implied that the visual system encodes such contextual features even under suppression (Roe, Lu, & Hung, 2005). Experiment 3 directly tested whether participants could unconsciously discriminate between physically bright stimuli and those made brighter through the illusion alone. Participants performed above chance in differentiating the two, which indicated that although subjective enhancements do not facilitate faster access to awareness, they are nonetheless processed to a meaningful extent beneath the threshold of consciousness (Harris et al., 2011).

These results raise an important question: if the visual system can unconsciously register illusory brightness, why does this not influence the speed at which suppressed stimuli reach awareness? The dissociation between detection and breakthrough implies that unconscious processing is not monolithic but consists of multiple stages, some of which may not impact conscious access directly. Prior research suggests that while mid-level features, such as symmetry, curvature, or local contrast, can be processed without awareness (Mudrik et al., 2011), only specific low-level signals—especially those encoded early in the visual hierarchy—can consistently survive under CFS (Yuval-Greenberg & Heeger, 2013). This suggests that not all unconscious representations are equally weighted in the competition for awareness and that a framework that distinguishes between early sensory encoding and later contextual integration may be necessary to account for the current findings.

We propose a two-tiered model of unconscious visual processing to interpret these findings. First, physical luminance appears to dominate the initial competition for awareness, which aligns with previous findings that higher contrast or luminance leads to faster emergence under CFS (Fang & He, 2005; Stein et al., 2011). From an evolutionary perspective, such a bias makes sense: physically intense signals (e.g., a flash of light) often convey urgent environmental changes and thus demand rapid processing (Gayet, Van der Stigchel, & Paffen, 2014). On a neural level, actual luminance likely produces stronger, more reliable activity in the primary visual cortex (V1), which gives these stimuli a competitive edge in interocular rivalry (Song & Yao, 2016). Second, perceptual brightness illusions—such as the glare effect—are likely encoded at mid-level or higher-level stages of the visual hierarchy and are often associated with areas such as V2 or V4 (Grossberg & Todorovic, 1988; Roe et al., 2005). These areas appear capable of contextual integration even without awareness, as shown in Experiments 2 and 3. However, the illusion’s failure to accelerate breakthrough suggests that unconscious representations of such effects remain weaker or more localized than those of physically brighter stimuli. One possibility is that while V2 may compute surface brightness gradients, the gating function of V1 in early interocular competition remains relatively impervious to top-down or mid-level feedback (Moors, Hesselmann, Wagemans, & Ee, 2017). As a result, the visual system can unconsciously register the illusion without sufficient low-level weight to impact BT. This aligns with hierarchical theories suggesting that conscious perception depends on the effective propagation of signals through both early sensory and higher integrative areas (Sterzer, Stein, Ludwig, Rothkirch, & Hesselmann, 2014).

These findings also echo broader evidence that unconscious vision can integrate certain contextual and surface-based illusions, even if this does not lead to conscious awareness (Harris et al., 2011). For example, Harris and colleagues demonstrated that simultaneous brightness contrast survives under CFS, albeit in attenuated form. Similarly, surface-based illusions involving local properties, such as color or contrast, can persist to some extent during suppression (Maruya, Watanabe, & Watanabe, 2008). Our results extend these findings by showing that even a more complex radial brightness illusion, such as the glare effect, is encoded unconsciously, as evidenced by above-chance performance in forced-choice discrimination (Experiment 3). In contrast, illusions that require global completion—such as Kanizsa figures—typically fail to manifest if their inducers are suppressed (Harris et al., 2011). This distinction suggests that illusions vary in their dependence on conscious processing. As a surface-based phenomenon, the glare illusion seems to rely on partial contextual interpolation rather than full contour formation. This aligns with models positing that mid-level mechanisms (e.g., V2 or V4) can operate in the absence of awareness, whereas global object-based illusions likely depend on more elaborate, feedback-driven assembly across multiple regions (Sterzer et al., 2014).

In sum, our study supports a nuanced view of unconscious vision. Illusions with low global demands—such as local brightness gradients—can be encoded without awareness, which allows the visual system to register meaningful contextual information even under suppression. However, these illusions lack the neural salience to compete with physically strong stimuli for conscious access. This dual outcome suggests that unconscious perception is neither minimal nor all-encompassing: it can extract substantial mid-level structure but remains bound by the architecture and priority rules of the visual hierarchy. Future work might explore how factors such as attention, familiarity, or prior learning influence these unconscious computations, particularly in terms of whether higher-level feedback can modify low-level visual competition under suppression.

## Conclusion

Our three experiments collectively provide two key insights into unconscious brightness processing. First, illusions do not act like real light intensity in accelerating access to awareness: physical luminance remains the key determiner of BT. Second, contextual brightness illusions are not simply lost below awareness—they can be discriminated against under forced-choice conditions, which indicates that mid-level or secondary visual areas register illusory brightness during suppression. Future research could investigate whether the manipulation of top-down attention or the engagement of different cortical pathways might allow illusions to exert more influence on BTs or whether illusions of color, motion, or depth behave differently from those rooted purely in brightness gradients. The employment of neuroimaging approaches (fMRI, EEG, or TMS) could help pinpoint precisely which brain regions mediate these unconscious brightness effects and shed further light on how and when conscious vision emerges from the interplay of low-level signals and higher-level contextual constructions.

## References

Adelson, Edward H. (n.d.). 24 lightness perception and lightness illusions. Retrieved from https://api.semanticscholar.org/CorpusID:17847203

Adelson, Edward H. (1993). Perceptual organization and the judgment of brightness. Science, 262(5142), 2042–2044.

Agostini, T., & Galmonte, A. (2002). A new effect of luminance gradient on achromatic simultaneous contrast. Psychonomic Bulletin & Review, 9(2), 264–269.

Blakeslee, B., & McCourt, M. E. (1999). A multiscale spatial filtering account of the white effect, simultaneous brightness contrast and grating induction. Vision Research, 39(26), 4361–4377.

Brainard, D. H., & Vision, S. (1997). The psychophysics toolbox. Spatial Vision, 10(4), 433–436.

Fang, F., & He, S. (2005). Cortical responses to invisible objects in the human dorsal and ventral pathways. Nature Neuroscience, 8(10), 1380–1385.

Faul, F., Erdfelder, E., Lang, A.-G., & Buchner, A. (2007). G* power 3: A flexible statistical power analysis program for the social, behavioral, and biomedical sciences. Behavior Research Methods, 39(2), 175–191.

Gayet, S., Van der Stigchel, S., & Paffen, C. L. (2014). Breaking continuous flash suppression: Competing for consciousness on the pre-semantic battlefield. Frontiers in Psychology, 5, 460.

Grossberg, S., & Todorovic, D. (1988). Neural dynamics of 1-D and 2-D brightness perception: A unified model of classical and recent phenomena. Perception & Psychophysics, 43(3), 241–277. 10.3758/bf03207869

Harris, J. J., Schwarzkopf, D. S., Song, C., Bahrami, B., & Rees, G. (2011). Contextual illusions reveal the limit of unconscious visual processing. Psychological Science, 22(3), 399–405.

Jiang, Y., Costello, P., & He, S. (2007). Processing of invisible stimuli: Advantage of upright faces and recognizable words in overcoming interocular suppression. Psychological Science, 18(4), 349–355.

Kingdom, F. A. (2003). Levels of brightness perception. In Levels of perception (pp. 23–46). Springer.

Kinzuka, Y., Sato, F., Minami, T., & Nakauchi, S. (2021). Effect of glare illusion-induced perceptual brightness on temporal perception. Psychophysiology, 58(9), e13851.

Mario, K., David, B., Denis, P., Allen, I., Richard, M., Christopher, B., et al. (2007). What’s new in psychtoolbox-3. Perception, 36(14), 1–1.

Martinsen, M. M., Kinzuka, Y., Sato, F., Minami, T., & Nakauchi, S. (2023). Breakthrough time depends on letter type and upright orientation–a pilot study using continuous flash suppression–. International Journal of Affective Engineering, 22(2), 157–165.

Martinsen, M. M., Yoshino, K., Kinzuka, Y., Sato, F., Tamura, H., Minami, T., & Nakauchi, S. (2024). Facial ambiguity and perception: How face-likeness affects breaking time in continuous flash suppression. Journal of Vision, 24(9), 18–18.

Maruya, K., Watanabe, H., & Watanabe, M. (2008). Adaptation to invisible motion results in low-level but not high-level aftereffects. Journal of Vision, 8(11), 7–7.

Moors, P., Hesselmann, G., Wagemans, J., & Ee, R. van. (2017). Continuous flash suppression: Stimulus fractionation rather than integration. Trends in Cognitive Sciences, 21(10), 719–721.

Mudrik, L., Breska, A., Lamy, D., & Deouell, L. Y. (2011). Integration without awareness: Expanding the limits of unconscious processing. Psychological Science, 22(6), 764–770.

Nuutinen, M., Mustonen, T., & Häkkinen, J. (2018). CFS MATLAB toolbox: An experiment builder for continuous flash suppression (CFS) task. Behavior Research Methods, 50(5), 1933–1942.

Paradiso, M. A., & Nakayama, K. (1991). Brightness perception and filling-in. Vision Research, 31(7-8), 1221–1236.

Pelli, D. G., & Vision, S. (1997). The VideoToolbox software for visual psychophysics: Transforming numbers into movies. Spatial Vision, 10, 437–442.

Reid Jr, R. C., & Shapley, R. (1988). Brightness induction by local contrast and the spatial dependence of assimilation. Vision Research, 28(1), 115–132.

Roe, A. W., Lu, H. D., & Hung, C. P. (2005). Cortical processing of a brightness illusion. Proceedings of the National Academy of Sciences, 102(10), 3869–3874.

Shapley, R., & Reid, R. C. (1985). Contrast and assimilation in the perception of brightness. Proceedings of the National Academy of Sciences, 82(17), 5983–5986.

Sinha, P., Crucilla, S., Gandhi, T., Rose, D., Singh, A., Ganesh, S., … Bex, P. (2020). Mechanisms underlying simultaneous brightness contrast: Early and innate. Vision Research, 173, 41–49.

Song, C., & Yao, H. (2016). Unconscious processing of invisible visual stimuli. Scientific Reports, 6(1), 38917.

Stein, T., Hebart, M. N., & Sterzer, P. (2011). Breaking continuous flash suppression: A new measure of unconscious processing during interocular suppression? Frontiers in Human Neuroscience, 5, 167.

Sterzer, P., Stein, T., Ludwig, K., Rothkirch, M., & Hesselmann, G. (2014). Neural processing of visual information under interocular suppression: A critical review. Frontiers in Psychology, 5, 453.

Suzuki, Y., Minami, T., Laeng, B., & Nakauchi, S. (2019). Colorful glares: Effects of colors on brightness illusions measured with pupillometry. Acta Psychologica, 198, 102882.

Tamura, H., Nakauchi, S., & Koida, K. (2016). Robust brightness enhancement across a luminance range of the glare illusion. Journal of Vision, 16(1), 10. 10.1167/16.1.10

Tsuchiya, N., & Koch, C. (2005). Continuous flash suppression reduces negative afterimages. Nature Neuroscience, 8(8), 1096–1101.

Willenbockel, V., Sadr, J., Fiset, D., Horne, G. O., Gosselin, F., & Tanaka, J. W. (2010). The SHINE toolbox for controlling low-level image properties. Hist, 1, 2.

Williams, S. M., McCoy, A. N., & Purves, D. (1998). An empirical explanation of brightness. Proceedings of the National Academy of Sciences, 95(22), 13301–13306.

Yang, E., Blake, R., & McDonald, J. E. (2010). A new interocular suppression technique for measuring sensory eye dominance. Investigative Ophthalmology & Visual Science, 51(1), 588–593.

Yoshida, A., Mittner, M., Mantiuk, R., & Seidel, H.-P. (2008). Brightness of glare illusion. Creem-Regehr, Sarah; Myszkowski, Karol: Symposium on Applied Perception in Graphics and Visualization : Proceedings APGV 2008, ACM, 83-90 (2008), 83–90. 10.1145/1394281.1394297

Yuval-Greenberg, S., & Heeger, D. J. (2013). Continuous flash suppression modulates cortical activity in early visual cortex. Journal of Neuroscience, 33(23), 9635–9643.

Zavagno, D. (1999). Some New Luminance-Gradient Effects. Perception, 28(7), 835–838. 10.1068/p2633

Zavagno, D., & Caputo, G. (2001). The glare effect and the perception of luminosity. Perception, 30(2), 209–222.

